# Proteotyping reactivant toxoplasmic encephalitis reveals virulence-associated dense granule protein GRA5 polymorphisms

**DOI:** 10.64898/2026.02.01.702898

**Authors:** R.A. Montoro, B. Chadwick, C. Su, K. Overmyer, J. Coon, L.J. Knoll, R. Striker

**Affiliations:** University of Wisconsin – Madison; University of Tennessee, Knoxville

## Abstract

A fatal case of toxoplasmic encephalitis, and others like it, has caused microbiologists and clinicians to question whether different strains of *T. gondii* have more pathogenic potential than others. This raises significant concern, as *T. gondii* is a widely spread parasitic organism that is presumed to lie dormant in a third of the world’s general population. In this study, we expand on a previously published proteomic analysis reactivated toxoplasmic encephalitis and have been able to identify *T. gondii*-specific peptides in the cerebrospinal fluid (CSF) of this patient and two additional cases of toxoplasmic reactivation. Multilocus PCR-restriction fragment-length polymorphism (PCR-RFLP) was used to genetically identify the *T. gondii* strain that resulted in this fatal case, which belonged to ToxoDB PCR-RFLP genotype #7. Using other *T. gondii* strains of the same genotype, we performed bioassays to compare the pathogenicity of this genotype with that of a clinically relevant strain, ME49. Both of the tested genotype #7 strains appear to have a greater pathogenic potential, although through likely different mechanisms. Of the most abundant *T. gondii-*specific CSF peptides across multiple patients, we identified a polymorphic region of the dense granule protein GRA5 that appears to have strain specificity. This approach could represent a “*proteotype”* that allows for *T. gondii* strain risk stratification within clinical samples. Ultimately spinal fluid could be a valuable tool in distinguishing between T. gondii exposed individuals with no cyst forms in the brain versus those exposed individuals that harbor “clinically silent” but viable brain cysts.

**SIGNIFICANCE:** A third of the world’s population is exposed to the parasite *Toxoplasma gondii*, residing in a dormant, encysted stage within neurons. *T. gondii* is a diverse microorganism with some strains having greater reactivation potential. There is no means of identifying which individuals are at risk of reactivation. Proteomic analysis of cerebrospinal fluid from patients with reactivated toxoplasmosis demonstrated a consistent pattern of *T. gondii* peptides, including GRA5, a secreted virulence factor. Postmortem analysis identified a ToxoDB PCR-RFLP genotype #7 strain associated with a fatality. Cerebrospinal fluid could provide clues to persistent brain infection and may identify strains able to reactivate.

## INTRODUCTION

*Toxoplasma gondii* is arguably the most successful parasitic organism to have evolved. It is theoretically capable of infecting any warm-blooded animal cell.^1^ While the feline is the definitive host, wherein sexual reproduction strictly occurs within the feline gut,^2^ infection of other animals results in the parasite replicating asexually and forming tissue cysts, mostly in neural and striated muscle tissues. Consumption of contaminated produce or infected intermediate hosts (*e.g*., livestock) is the primary route of infection in humans,^3^ and subsequently, a third of humanity has been exposed to *T. gondii*.^4^ While an acute infection and subsequent systemic dissemination of *T. gondii* in an immunocompetent host often presents asymptomatically or as mild flu-like symptoms,^5^ not all strains behave uniformly.^6–8^

Amidst the rising AIDS epidemic in the 1990s, *T. gondii* was recognized as a major AIDS opportunistic infection. Research suggested that there were only three major lineages and relatively little phenotypic diversity between lineages.^9,10^ However, subsequent analysis using multilocus polymerase-chain reaction restriction-fragment length polymorphism (PCR-RFLP) and multilocus microsatellite genotyping approaches, and later whole genome sequencing, now suggests a much more varied and complex reality.^11–14^

Due to resource and time constraints, much of the data on *T. gondii* focuses on the acute infectious stage and on describing differences in pathogenicity among strains during earlier phases of infection. While the chronic stage of *T. gondii* receives less attention, it carries substantial health risks. Immunocompromised people are at significant risk for reactivation of *T. gondii*. For example, HIV positive individuals with a CD4 count <100 cells/μL have a 30% risk for reactivation.^15^ Toxoplasmic encephalitis (TE), a potential disease incurred from reactivation, can result in a myriad of symptoms. TE can present subtly as a headache and confusion, or more aggressively as focal neural deficits, seizure, or even coma.^16^ While this increase in risk is proportionally higher in HIV positive populations,^15^ the incidence of reactivation is likely underreported due to its often subtle onset. Within humans, this chronic infection is presumed to reside throughout the entirety of the host’s lifespan. Therefore, improving our ability to distinguish between strains that have greater pathogenic capability is of great importance.

We previously published a short report on a reactivation case that, unfortunately, resulted in a fatal outcome at our institution in 2023. This case and others like it provide evidence to inform healthcare practitioners of the risk that starting anti-TNF-α therapy in individuals with chronic *T. gondii* infections may pose. However, at the time, we were unable to identify the strain that resulted in this fatal case. Additionally, we presented a proteomic analysis of the patient’s cerebrospinal fluid.^17^ This was done in an effort to provide proof of concept for a proteomic approach toward identifying *T. gondii* strains, a *“proteotype*”.

In this study, we continue our follow-up investigation of the *T. gondii* strain that resulted in a fatal toxoplasmic encephalitis. We expanded our investigation into the CSF proteomic analysis of the aforementioned case and 2 other individuals who were also diagnosed with TE, who fortunately have recovered. We pursued genetic characterization using RFLP genotyping of the fatal case and followed up with a phenotypic characterization by performing *in vivo* experimentation of strains within the same RFLP genotype to investigate their pathogenicity compared to a clinically relevant strain of *T. gondii*, ME49. Lastly, we identified a recurring peptide across all patients from the proteomic analysis, GRA5, with polymorphisms that correlate with the variation of genotype #7 mouse virulence. This extends prior mouse data (CITE) that identifies GRA5 as a virulence factor to human data.

## METHODOLOGY

### Restriction Fragment Length Polymorphism Genotyping

In agreement with the patient’s family, an autopsy was performed at the UW University Hospital. Multiple brain autopsy specimens were collected for multilocus PCR-RFLP genotyping.

Twenty-five mg each of five brain tissue sections were taken for DNA extraction using DNeasy Blood and Tissue kit (Qiagen; Cat No. 69504) following the manufacturer’s instructions. DNA was resuspended in 50 μL of buffer AE and then subjected to PCR-RFLP analysis using 10 genetic markers (SAG1, SAG2, SAG3, BTUB, GRA6, c22-8, c29-2, L358, PK1, and Apico).^11^

### Parasite Maintenance

Three strains of *T. gondii* were used for *in vivo* experimentation. Genotype #7 strains were graciously provided by the Su lab; G622M and TgCkBr112 were previously isolated from chronically infected animals. ^9,18^ Type II strains predominate clinical cases in North America;^9^ thus, the Type II, ME49 strain served as a clinically relevant control. All strains were maintained and passaged in human foreskin fibroblast cell cultures before intraperitoneal infection.

### Mouse Infections and Sickness Score

A total of 60 house-bred female C57Bl6/J mice were used to assess the virulence of the three *T. gondii* strains. All mice were treated according to the guidelines established by the Institutional Animal Care and Use Committee (IACUC) of the University of Wisconsin School of Medicine and Public Health (protocol number M005217). The institution adheres to the regulations and guidelines set by the National Research Council. Animals were intraperitoneally infected with either [2×10^1^] or [2×10^4^] parasites (N=10 per group) and monitored daily for behavioral abnormalities. A 5-point sickness score was used to evaluate animal health in these experiments, which assessed for the following behaviors: lack of grooming, eye squinting, nose bulging, hunching, and complete lack of movement. Animals were euthanized via CO_2_ if a total of 5 points was recorded on any day.

### Brain Processing, Staining, Cyst Counts, and Serology

At the experimental endpoint, 28 days post-infection (dpi), all surviving animals were euthanized. Blood was collected, serum was separated, and immediately stored in -80°C for later use. Brains were dissected and stored in PBS at 4°C overnight. The following day, brains were homogenized and fixed in 3% formaldehyde. Homogenized tissue was stained using a fluorescein-conjugated dilochos biflorus agglutinated stain (Vector Laboratories, Catolog # FL-1031). Aliquots (15 μL) of processed brains were mounted on glass coverslips and blinded. Cyst quantification was performed using a Zeiss Axioplan III microscope at 10X. Animals without evidence of cysts had the rest of their homogenized brain tissue remounted (80 μL+) and inspected for the presence of at least one tissue cyst. GraphPad PRISM software was used to visualize and perform statistical analyses.

### Sample Preparation for Proteomics

Samples were prepared as previously described.^17,19^ The CSF sample was brought to 90% methanol and incubated with rocking at 4°C for 30 minutes. The sample was then centrifuged at 4°C for 5 minutes at 14,000 x g and the methanol removed. The precipitate was resolubilized in lysis buffer (8 M urea, 10 mM TCEP, 40 mM CAA, 100 mM Tris) and protein concentration determined by a Pierce BCA assay (Thermo Fisher). The sample was then diluted with 100mM Tris so the final concentration of urea was 1.5M, and 50µg of protein was digested with Trypsin (Promega) and Lys-C (Sigma) in a 50:1 ratio of protein/enzyme overnight on a rocker at room temperature. Enzyme activity was quenched by adding trifluoroacetic acid (TFA) to a final pH of 2.0. The peptides were then desalted using 10mg Strata-X Polymeric Solid Phase Extraction cartridges (Phenomenex) and eluted in 300µl of elution buffer (80% ACN, 0.2% TFA). The peptides were then dried with the SpeedVac before proteome analysis.

### Liquid Chromatography-Mass Spectrometry (LC-MS) Analysis and Proteomic Analysis

Peptide samples were analyzed using a Vanquish Neo UHPLC system (Thermo Scientific) coupled to an Orbitrap Astral mass spectrometer (Thermo Scientific) via a Nanospray Flex ionization source (Thermo Scientific). A 40 cm fused silica capillary column (75 µm i.d., 360 µm o.d.; Polymicro Technologies) was pulled, etched, and packed in-house with 1.7 µm C18 particles (Waters), as previously described.^20^ 500ng of peptides were injected and separated at 55 °C using a 30-minute active gradient at a flow rate of 300 nL/min. Mobile phase A consisted of 0.2% formic acid in water, and mobile phase B consisted of 0.2% formic acid in 80% acetonitrile. The gradient ramped from 0% to 4% B over the first 2 minutes, followed by an increase to 45% B by 30 minutes using a pump curve setting of 5. Data were acquired in data-independent acquisition (DIA) mode. Full MS1 scans were collected in the Orbitrap at a resolution of 240,000 across a mass range of 380–980 m/z, with a normalized AGC target of 250%, a maximum injection time of 50 ms, and an RF lens setting of 40%. Astral MS2 scans were performed with a maximum injection time of 2.5 ms, a scan range of 150–2000 m/z, an RF lens setting of 30%, HCD collision energy of 30%, a normalized AGC target of 500%, and an isolation window of 2 m/z, resulting in 300 scan events per cycle.

Mass spectrometry data were processed using Spectronaut directDIA (version 19.7)^21^ with default settings and the *T. gondii* protein fasta datasets from ToxoDB version 68.^22^

### GRA5 PCR Amplification and Sequencing

Parasites maintained in cell culture were scraped and syringe-passaged, followed by DNA extraction (Zymo Research Quick-DNA Miniprep Plus Kit, Catalog No. D4068). These strains and the two patient-derived DNA samples used to identify the strain as genotype #7 were subjected to a nested PCR amplification using newly designed and validated primers (Supplementary Table 1, Supplementary Figure 1). Crude PCR amplified DNA and internal GRA5 primer pairs were sent to Functional Biosciences for Sanger sequencing of the amplified GRA5 region of interest. Sequence alignment and identification of nucleotide polymorphisms between these strains were performed using Geneious Prime software. ExPasy online software was used to translate the nucleotide sequence into the amino acid sequence for comparison.

## RESULTS

Three patients at the UW Health System were diagnosed with reactivated toxoplasmosis. One patient succumbed to the disease (denoted as EM), and the other two survived (ARA and WB). During their clinical workup, CSF was collected and assessed for infectious etiology. CSF was subjected to LC-MS, and peptide analysis was performed.

### CSF Proteomic Analysis

Proteomic analysis of CSF yielded multiple *T. gondii*-specific peptides consistently present across all three patients. Of note, 14-3-3 protein, dense granule protein GRA1, microneme protein MIC5, and dense granule protein GRA5 were identified as the top four most abundant peptides across all three samples. ToxoDB was used to provide a reference genome to identify these proteins as *T. gondii*-specific, in order to identify strains that had matching sequences to any of these proteins. Among these 4 identified peptides, fifteen, four, eight, and four (14-3-3 protein, GRA1, MIC5, and GRA5, respectively) *T. gondii* strains shared the same peptide sequence across all patients (Figure 1).

**Figure 1:**
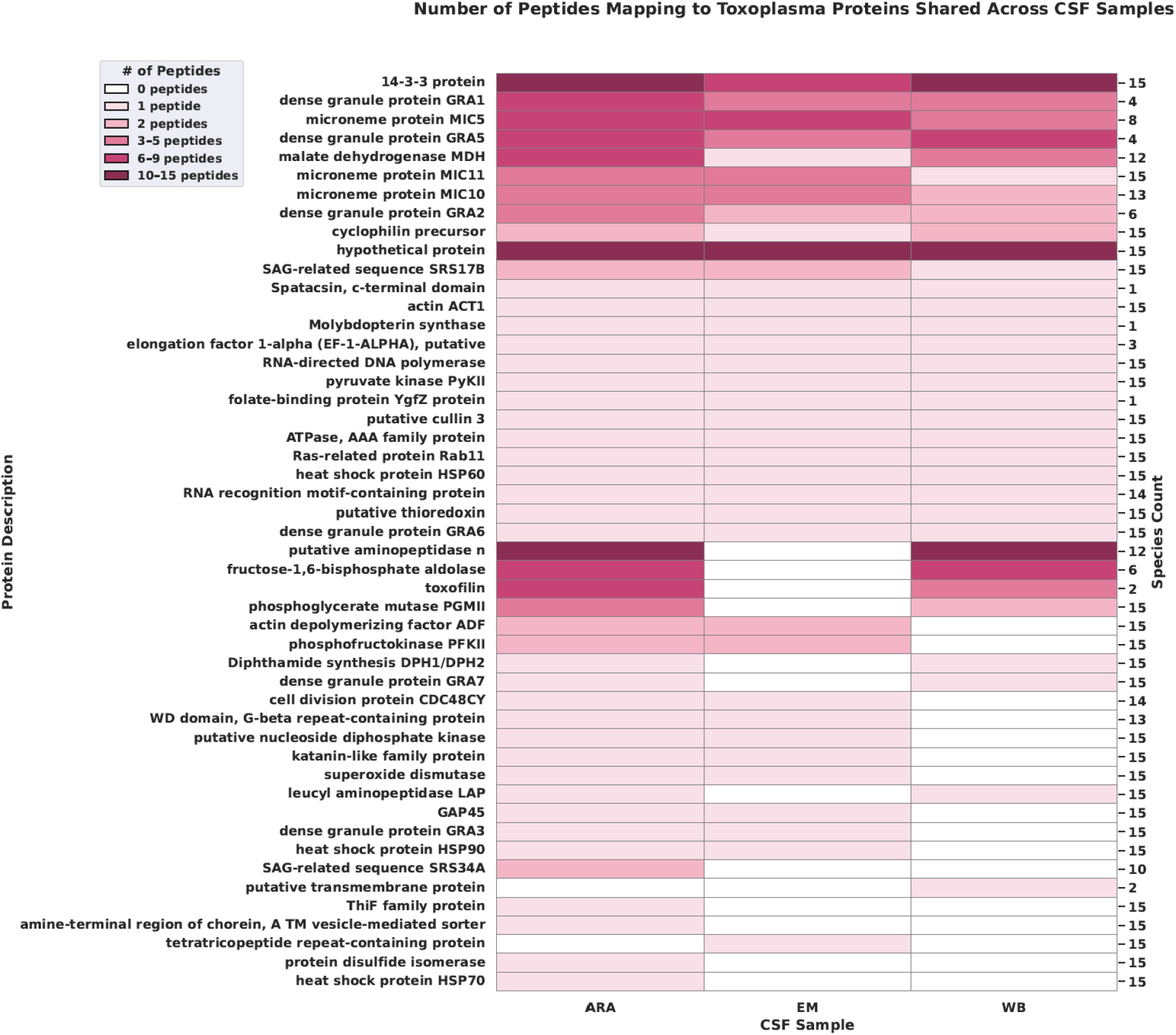
Proteomic Analysis of CSF from Patients with Toxoplasmic Encephalitis Identified Multiple Overlapping *T. gondii*-specific Peptides. -LC-MS and proteomic analysis of 3 distinct TE cases, 1 fatal and 2 nonfatal, yielded multiple overlapping *T. gondii*-specific peptides matching multiple *T. gondii* strains. The x-axis denotes the three patients who had TE, with EM being the patient from the fatal case. Listed on the left y-axis is the description of the identified protein. The right y-axis denotes how many *T. gondii* strains were identified and consistently matched to one or more peptides across all three patients. The shading denotes the number of fragment peptides within each patient CSF sample. Four proteins; 14-3-3 protein, dense granule protein GRA1, microneme protein MIC5, and dense granule protein GRA5, yielded the highest number of fragment peptides across all three patient samples.

### RFLP Genotyping

DNA from two brain autopsy specimens from patient EM was isolated and genetically characterized using PCR-RFLP genotyping to identify them as genotype #7 (Table 1).

**Table 1.** RFLP Genotyping of Fatal Case Identified Strain as Genotype #7. -RFLP-PCR genotyping was implemented to identify the RFLP *T. gondii* genotype of the *T. gondii* strain that resulted in a fatal toxoplasmic encephalitis. Per this assay, it was confirmed from 9 out of 10 microsatellites that the strain resulting in this death belongs to genotype #7.

| Sample ID | SAG1 | (3' +5')<br>SAG2 | Alt<br>SAG2 | SAG3 | BTUB | GRA6 | C22-8 | C29-2 | L358 | PK1 | Apico | ToxoDB<br>PCR-RFLP<br>Genotype |
| --- | --- | --- | --- | --- | --- | --- | --- | --- | --- | --- | --- | --- |
| GT1<br>(reference) | I | I | I | I | I | I | I | I | I | I | I | # 10 |
| PTG<br>(reference) | II | II | II | II | II | II | II | II | II | II | II | # 1 |
| CTG<br>(reference) | II/III | III | III | III | III | III | III | III | III | III | III | # 2 |
| TgCgCa1<br>(reference) | I | II | II | III | II | II | II | u-1 | I | u-2 | I | # 66 |
| MAS<br>(reference) | u-1 | I | II | III | III | III | u-1 | I | I | III | I | # 17 |
| TgCatBr64<br>(reference) | I | III | III | III | III | III | I | I | I | u-1 | I | # 19 |
| TgCatBr64<br>(reference) | I | III | u-1 | III | III | III | u-1 | I | III | III | I | # 111 |
| TgRsCr1<br>(reference) | u-1 | I | II | III | I | III | u-2 | I | I | III | I | # 52 |
| Brain<br>(sample 1) | I | III | III | III | III | III | III | III | III | III | nd | # 7 |
| Brain<br>(sample 2) | I | III | III | III | III | III | III | III | III | III | nd | # 7 |

### In Vivo Bioassays

To study the pathogenic potential of genotype #7 strains, two additional genotype #7 strains available to the Su lab, G622M and TgCkBr112, were used.

Animals infected intraperitoneally with either [2×10^1^] or [2×10^4^] parasites were monitored daily for abnormal behavioral changes or death secondary to infection. 100% of animals infected with high-dose G622M succumbed to infection or were euthanized by 10 dpi. 80% of animals infected with low-dose G622M succumb to infection by 14 dpi. 100% of animals infected with high and low doses of ME49 and TgCkBr112 strains survived until experiment endpoint (Figure 2A).

**Figure 2:**
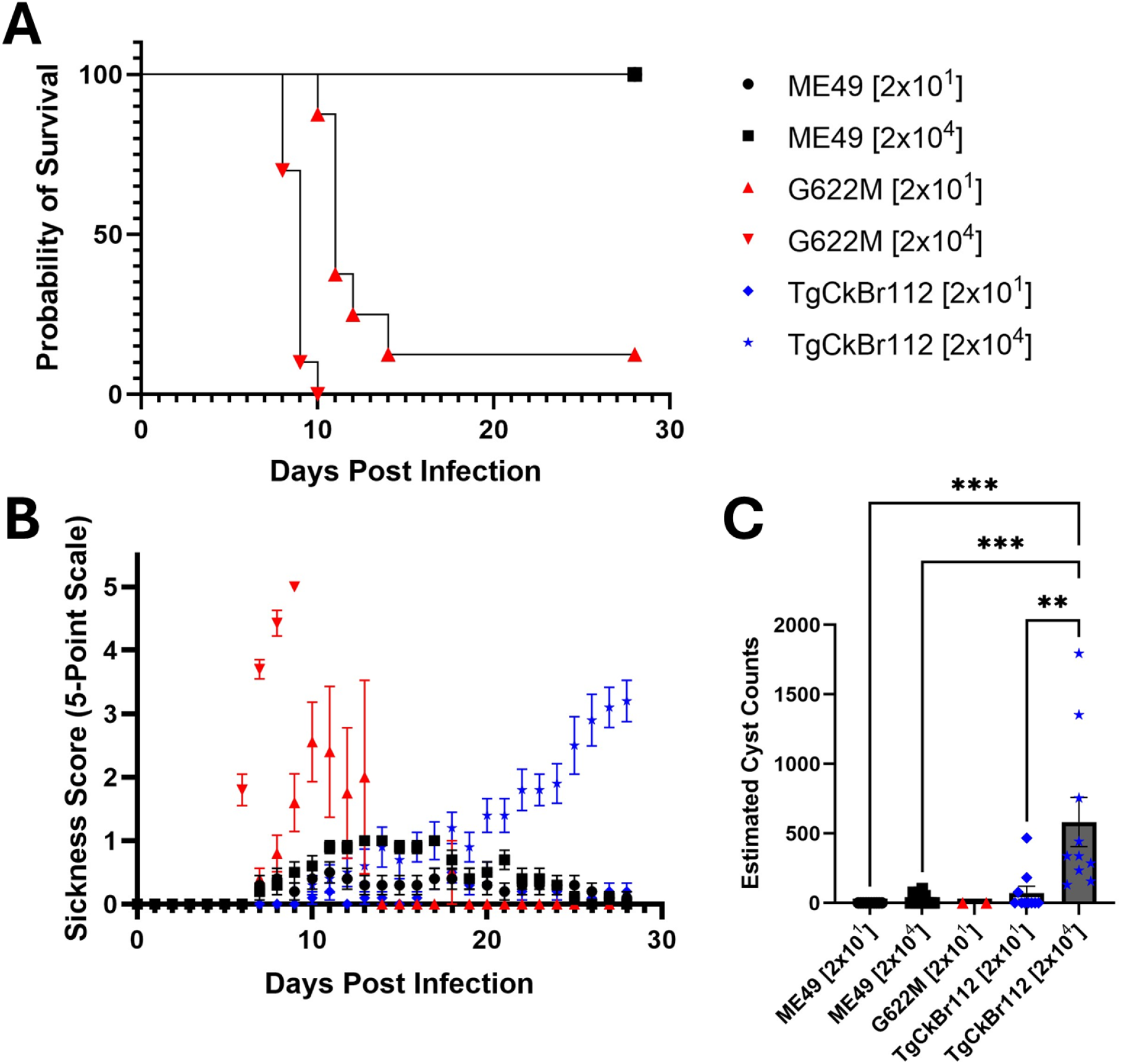
*In Vivo* Assays Demonstrate Genotype #7 Strains Have Greater Pathogenic Potential Compared to a Clinically Relevant Strain. -Animals were monitored daily on a 5-point sickness score scale. Animals with a 5/5 sickness score were immediately euthanized. The survival curve (Figure 2A) showed higher mortality during the experimental period in animals infected with the G622M strain than in those infected with ME49 or TgCkBr112. Sickness scores (Figure 2B) over the experimental period indicate a continuous increase in infection severity in animals infected with TgCkBr112 compared with ME49. Brain cyst count estimates (Figure 2C) were made at 28 dpi, and a 2-way ANOVA was performed to assess differences. The high dose of TgCkBr112 yielded the highest number of brain cysts compared to all other groups. Data shown as means +/-SEM.

Animals infected with high and low doses of ME49 had peak sickness scores at 14 and 11 dpi, with mean sickness scores <1. Sickness score of animals infected with high-dose G622M rose starting at 6 dpi and continued to increase until 9 dpi. Sickness score of animals infected with low-dose G622M began to rise at 7 dpi and peaked at a mean sickness score of 2.5 at 10 dpi, after which it began to fall. This decline in sickness score is due to the two surviving animals. Infection with high-dose TgCkBr112 resulted in a steadily rising sickness score at 9 dpi and continued to increase with a peak sickness score of 3 at 28 dpi. The low-dose TgCkBr112 resulted in much lower sickness score peaks of <1 at 11 dpi (Figure 2B).

Surviving animals were euthanized at 28 dpi to assess parasitic burden, assessed by brain cyst quantification. Animals infected with low-dose ME49 had on average <26 cysts per brain. The high-dose ME49 infection yielded an average 31.2 cysts per brain. Only 2 animals infected with either a high or low-dose G622M strain survived into the chronic stage. Animals infected with a low dose of G622M had <26 cysts per brain. The low dose of TgCkBr112 resulted in an average of 72.8 cysts per brain. The high dose of TgCkBr112 yielded an average of 582.8 cysts per brain by 28 dpi (Figure 2C). 2-way ANOVA of brain cyst counts revealed significant differences among groups (F(4,28) = 7.734, p = 0.0002). Post-hoc analysis revealed the high-dose TgCkBr112 yielded significantly more brain cysts than the low-dose ME49 (p = 0.0005), high-dose ME49 (p = 0.0009), and the low-dose TgCkBr112 (p = 0.0023). Low sample size for G622M group excluded it from analysis.

We did not observe the presence of cysts in several samples. For this assay, only 15 μL of homogenized brain tissue is used. To confirm infection occurred, the remaining homogenized tissue was mounted onto microscopy slides and inspected for the presence of any cysts (Supplementary Figure 2A). Animals without cysts in this follow-up assessment underwent serologic testing for anti-*T. gondii* antibodies in their serum (Supplementary Figure 2B) to confirm immune response to a *T. gondii* infection.

*T. gondii*-specific virulence factors, such as the rhoptry (ROPs) and dense granule proteins (GRAs), have garnered much attention.^23^ The top four peptides on the CSF analysis in this study included two GRA proteins, GRA1 and GRA5. Dense granule protein GRA5 is found within coccidian parasites, such as *T. gondii*, and plays a role in host-parasite interactions; notably, it manipulates host homeostasis to promote parasite survival. Importantly, unlike the other three top proteins, further investigation, through ToxoDB, of these proteins showed that GRA5 had no introns present in its genomic sequence, facilitating downstream sequencing and analysis of this protein.

### GRA5 Sequencing

Various fragment peptides from the proteomic analysis were matched to GRA5. Using BLASTp, certain peptide sequences were found to be exclusive to certain *T. gondii* strains (Supplementary Table 2). The GRA5 protein sequence is particularly interesting, as it contains multiple specific amino acid residues that differ between strains matched to the proteomic findings (Figure 3A).

**Figure 3:**
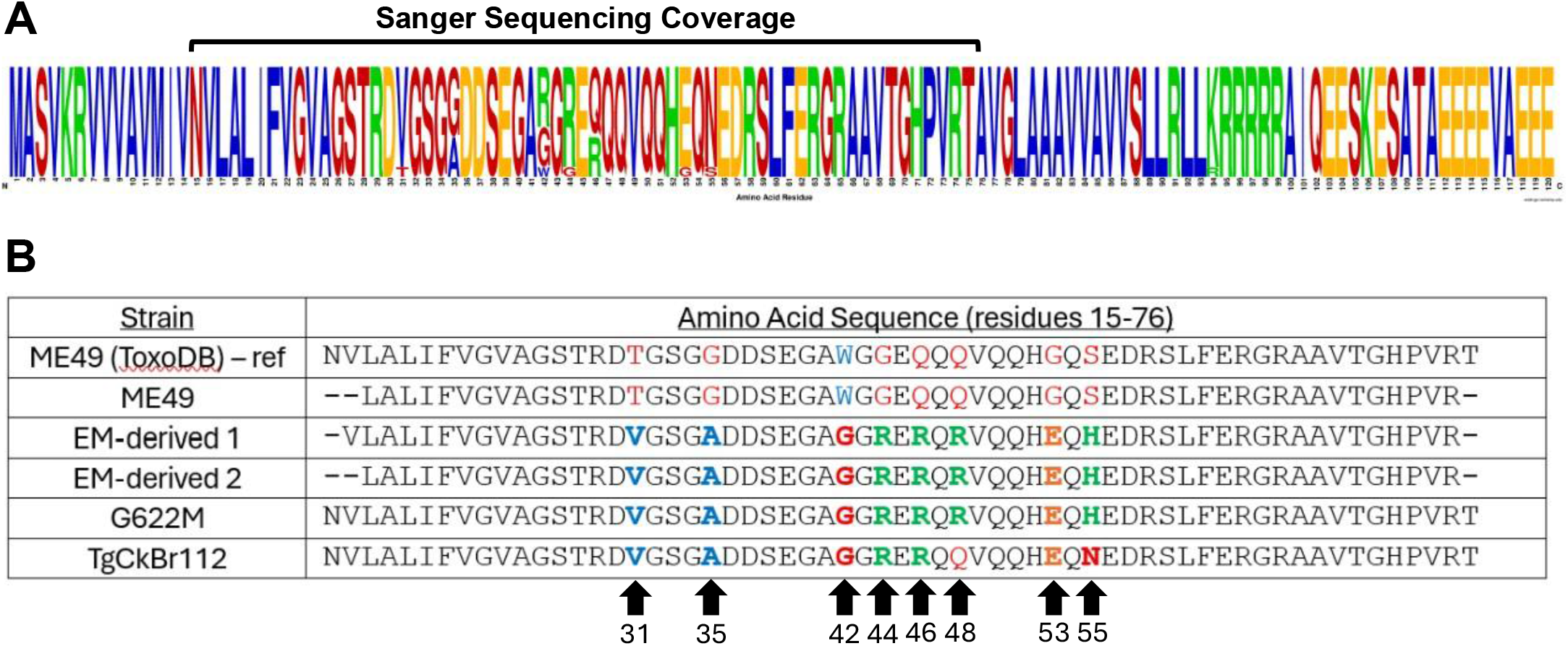
GRA5 Amino Acid Sequence Varies Across *T. gondii* Strains. -GRA5 amino acid sequence (available on ToxoDB) across all matching *T. gondii* strains identified on proteomic analysis allowed for the identification of multiple amino acid residues that differ between these strains (Figure 3A). Polymorphisms occurred at the amino acid positions 31, 35, 42, 44, 46, 53, 55, and 94 amino acid residues. The proteomic analysis covered all but the 94th amino acid residue. Amino acid sequence alignment among a reference sequence of ME49 has 100% homology with our cultured ME49 strain. With respect to the 58 overlapping amino acid residues, sequence homology between the three genotype #7 strains assessed shows 97% homology compared to 86-88% homology between genotype #7 strains and ME49 (Figure 3B).

Due to this variability in the dense granule protein GRA5 amino acid sequence, we sought to sequence the GRA5 protein from the fatal case of TE and assess if there was sequence homology across the other genotype #7 strains. Following DNA extraction and GRA5 nested PCR amplification (Supplemental Figure 1) of ME49, G622M, TgCkBr112, and two samples from EM, Sanger sequencing was performed to assess any polymorphic differences among these samples. gDNA sequence was aligned to an ME49 dense granule protein reference sequence obtained from ToxoDB (Supplementary Table 3). Translated amino acid sequence was aligned to the known ME49 GRA5 sequence obtained from ToxoDB (Figure 3B). It appears that genotype #7 strains have relatively conserved amino acid sequences. Of the 58 overlapping amino acid residues derived from Sanger sequencing, 86-88% homology is observed when comparing genotype #7 strains to ME49. Notably, an additional amino acid residue was identified to be polymorphic, residue 48, which had not been identified in the proteomic analysis. Six out of these eight variable amino acid residues were consistent between genotype #7 strains, with the TgCkBr112 strain differing from the other genotype #7 strains at residues 48 and 55. The amino acid residues differed significantly from the ME49 sequence in that at amino acid residues 31, 35, 42, 44, 46, 48, 53, and 55, genotype #7 strains had polymorphisms that resulted in amino acids with differing chemical properties from the ME49 reference sequence.

These data provide a foundation upon which CSF-derived proteomic analysis, with a specific focus on the dense granule protein GRA5, may offer sufficient resolution for identifying *T. gondii* strains that may have greater pathogenic potential.

## DISCUSSION

While chronic *T. gondii* infections are often asymptomatic, reactivation can occur, and there is evidence suggesting subtle neurologic effects from these persistent chronic cysts.^24,25^ Here, we provide evidence that the *T. gondii* strain associated with a fatal case of toxoplasmic encephalitis^17^ is a PCR-RFLP genotype #7 strain. After this genetic characterization, we tested if *T. gondii* strains identified as PCR-RFLP genotype #7 vary in pathogenic potential compared to ME49, a *T. gondii* strain considered clinically relevant with Europe and North America. This evidence suggests that some PCR-RFLP genotype #7 strains may be of greater clinical concern than previously thought.^18,26,27^

There is a clear difference in the pathogenesis of disease of the two genotype #7 strains assessed within *in vivo* models in this study. The G622M strain appears to induce significant mortality during the acute phase of the infection, whereas the TgCkBr112 strain appears to have a delayed onset of illness, at least within the C57Bl6/J mouse background. The TgCkBr112 strain results in a greater parasitic burden compared to ME49, suggesting a greater risk for reactivation in hosts infected with this parasite. The clear difference in disease progression suggests that these two PCR-RFLP genotype #7 strains may have different mechanisms through which they induce disease.

While the GRA5 polymorphism may not be entirely responsible for the phenotypic differences between the G622M and TgCkBr112 strains, it is not surprising that GRA5 is responsible for at least some of the phenotype as GRA5 has already been identified as a virulence factor (CITE Sibley GRA5 paper). Dense granule protein GRA5, specifically, is found among coccidian parasitic organisms. Within *T. gondii*, it plays a role in host invasion^28^ and parasite survival during its encysted stage^29^ in both secreted and transmembrane-bound actions. GRA5, along with other dense granule proteins (i.e., GRA3, GRA7, GRA8, and GRA14) is critical to forming the parasitophorous vacuole membrane and in this membrane appears to transition into a cyst membrane that surrounds the cyst wall.^30^

While GRA5 was found in all three analyzed CSF samples, other dense granule proteins and microneme proteins were also consistently present. The MIC5 protein has previously been recognized to encode the immunodominant antigen, H4,^31^ and has been shown to modulate the function of other invasion-associated *T. gondii* proteins, including a subtilisin protease responsible for activating parasite motility and invasion.^32^ More recent work has identified that GRA1 is crucial for parasite propagation during the acute stage of infection,^33^ and the *T. gondii* 14-3-3 protein has been proposed to increase host cell motility, facilitating systemic dissemination.^34^

In conclusion, while CSF proteomics is not needed to diagnose toxoplasmic encephalitis secondary to reactivation, it can provide important data to give some information on the strain of *T. gondii* resulting in disease. Even within PCR-RFLP genotype #7 strains, there exists phenotypic differences in at least mouse virulence. A number of chronic psychiatric and neurologic conditions have been associated with *T. gondii* but there are inconsistencies in the studies. More molecular information on which *T, gondii* strains people have been exposed to, and which strains may still be present in cerebral tissue could provide more clarity on possible clinical consequences of *T. gondii*. Given the genetic diversity of *T. gondii* and its high prevalence in the human population, particularly in South America, understanding the relationship of *T.gondii* genotypes and their potential in reactivation in chronic infection is important. Our study here clearly showed the variations of *T. gondii* “*proteotype”* ^35^ among the patients with different outcomes. Although our sample size is small, this study presents a novel tool for investigating reactivation of chronic toxoplasmosis in humans.

