## Supplemental Material for "Proteotyping reactivant toxoplasmic encephalitis reveals virulence-associated dense granule protein GRA5 polymorphisms"

**SUPPLEMENTARY MATERIAL**

**Supplementary Table 1:** Outer and internal nested PCR primer pair sequences used for GRA5 amplification and subsequent sequencing.

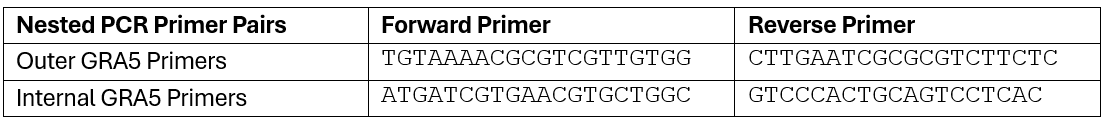

**Supplementary Figure 1:** Agarose gel electrophoresis of PCR amplified GRA5 gDNA.

- Validation of external and internal primer pairs was determined after PCR amplification (Supplemental Figure 1A) using the clinically relevant strain, ME49, and the 2 genotype #7 strains. Nested PCR was validated using the ME49 strain (Supplementary Figure 1B). Nested PCR in all samples was performed (Supplementary Figure 1C).

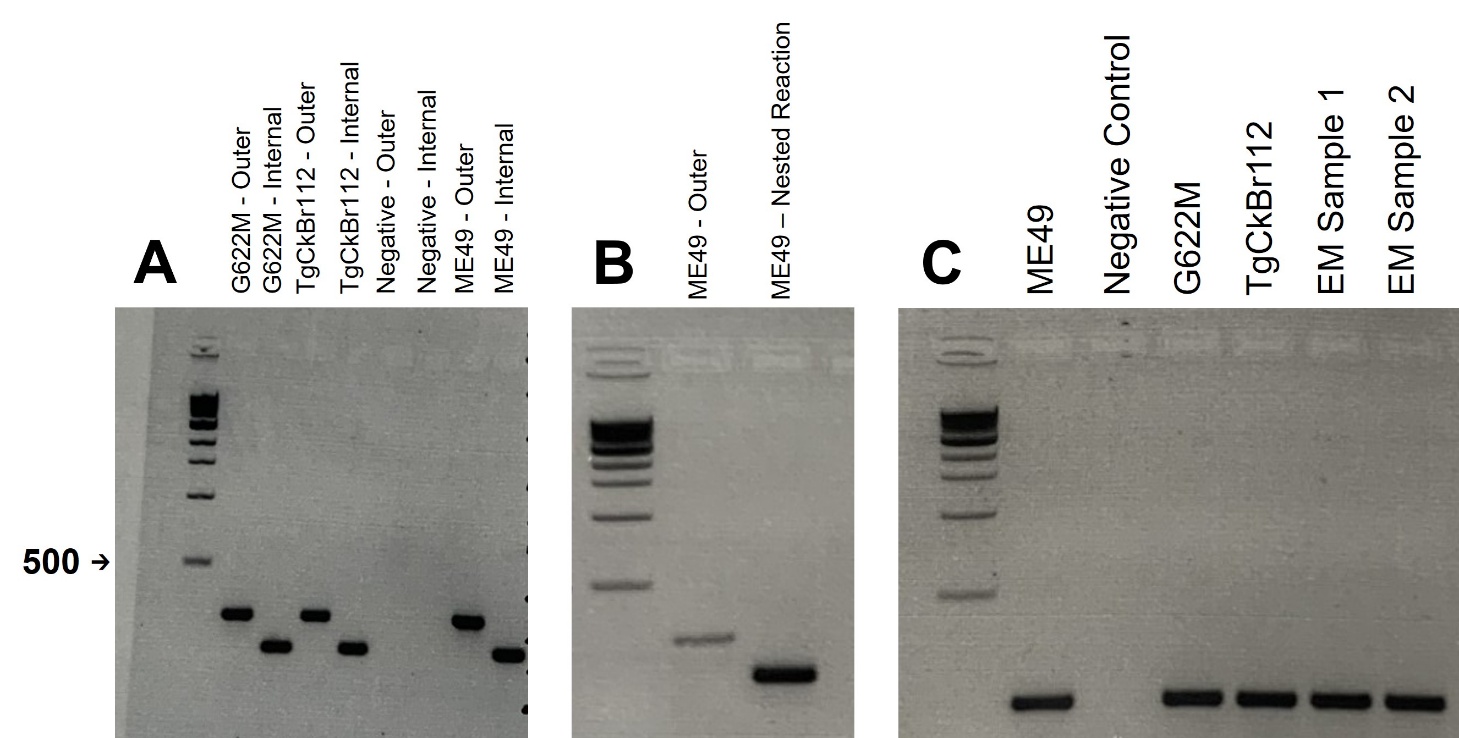

**Supplemental Figure 2:** Verification of infection in animals with without cysts.

- For animals without evidence of brain cysts with our initial assay, the remaining stained, homogenized brain tissue was mounted and scanned under a microscope for the presence of cysts. If a single cyst was identified, the animal was denoted to have at least one cyst, denoted as “Yes” in the table. All others that did not have cysts present were denoted as “No” (Supplemental Figure 2A). Whole ME49 parasites were lysed in radioimmunoprecipitation assay (RIPA) buffer. 100 μg of protein was separated on a 10% acrylamide gel and transferred to a nitrocellulose membrane. The membrane was cut into strips, blocked in PBS-Tween with 5% milk, and incubated with serum (1:1,000 in PBS-1% Tween). The strips were then incubated with a secondary antibody (1:5,000 goat anti-mouse HRP) and developed using ECL Prime Western Blotting Detection Reagent (Cytiva). The membrane was imaged on a Licor imager. All infected animals were revealed to have evidence of positive *T. gondii* serology despite no visual evidence for cysts in the brain (Supplementary Figure 2B).

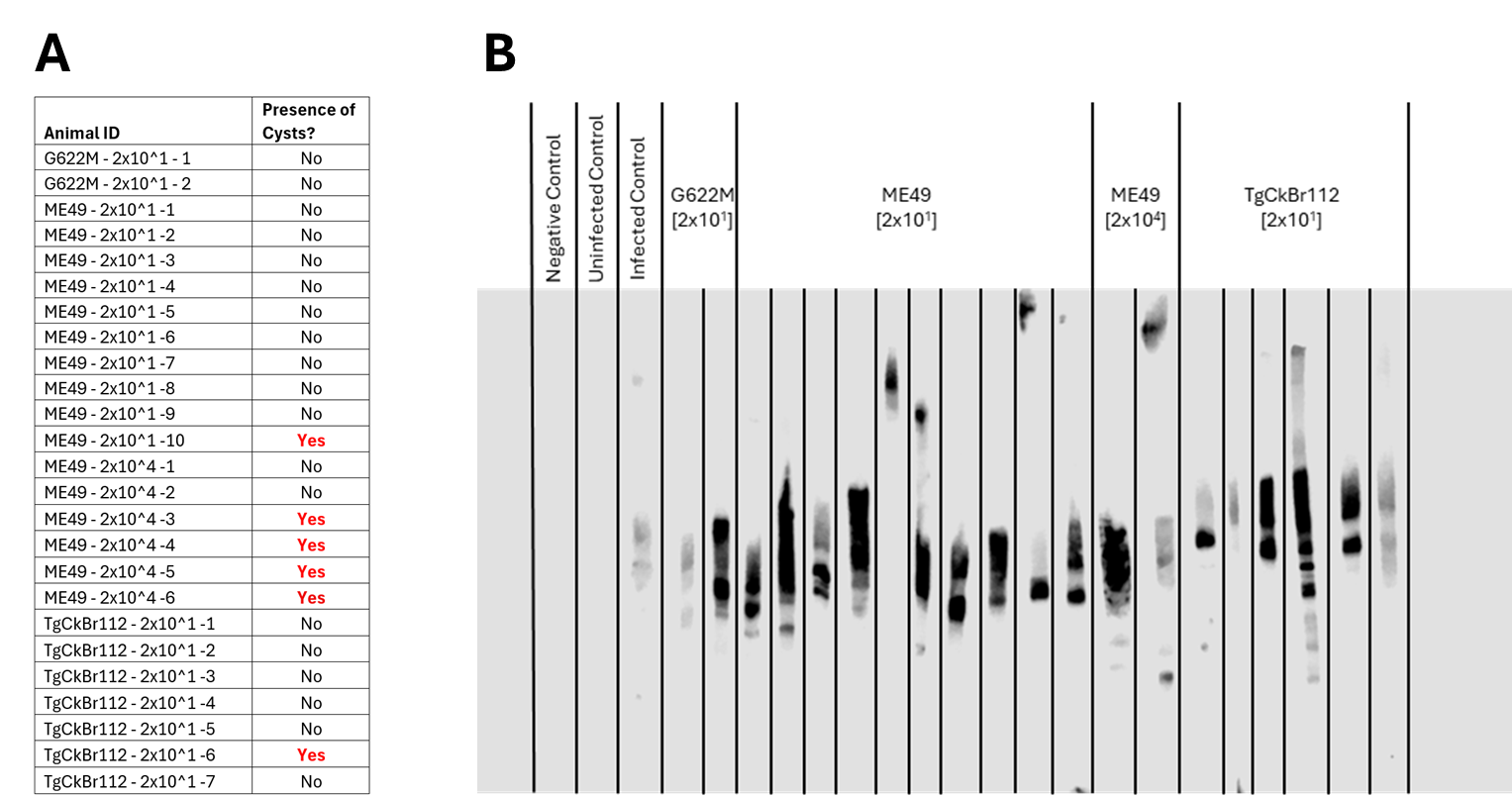

**Supplementary Table 2:** Proteomic Analysis Unveiled Multiple Peptide Sequences within the *T. gondii*-specific GRA5 Amino Acid Sequence Matching Several *T. gondii* Strains.

- Dense granule protein GRA5 yielded 9 fragment peptides. Among the identified peptide sequences, some were present in only two or all three patients, and two were unique to specific patients. These fragment peptides were cross-referenced using BLASTp, and strains with 100% homology are listed.

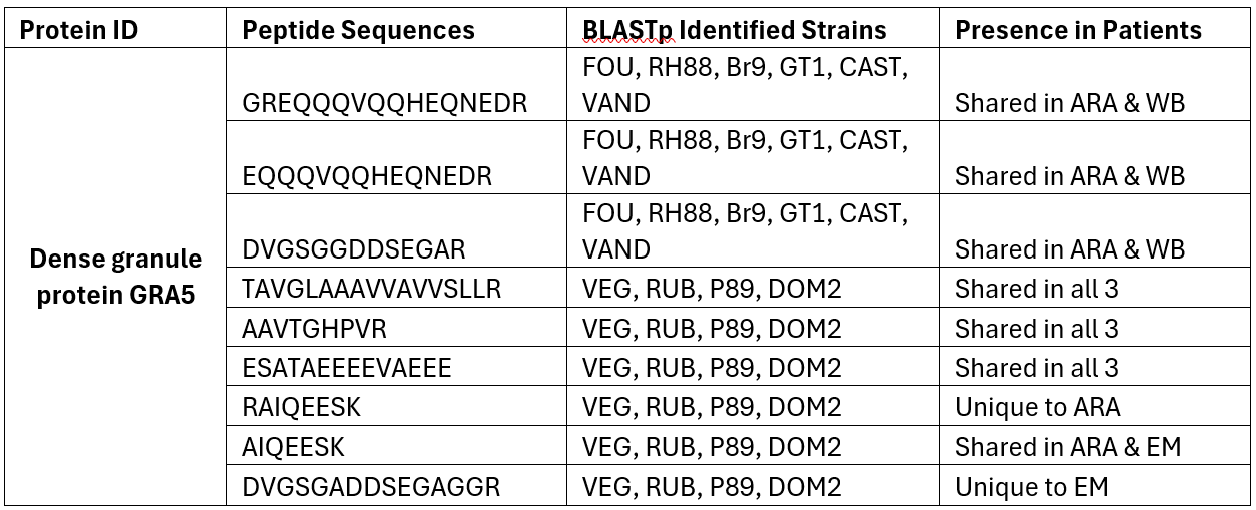

**Supplementary Table 3:** Sanger sequence nucleotide alignment.

- Within the overlapping sequenced region, nucleotide polymorphic sites were identified at 10 different sites, corresponding with the identified amino acid polymorphisms from the proteomic analysis.

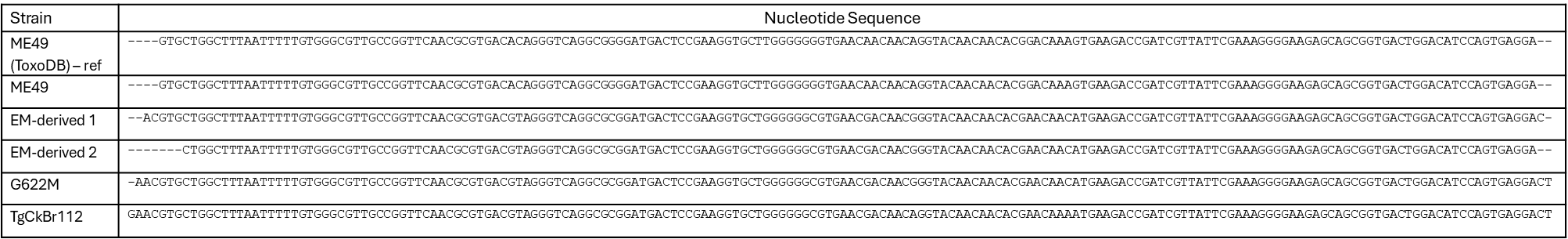
